# High throughput Bioluminescent assay to characterize and monitor the activity of SARS-CoV-2 Methyltransferases

**DOI:** 10.1101/2022.09.12.505486

**Authors:** Kevin Hsiao, Hicham Zegzouti, Said Goueli

## Abstract

The fast rate of viral mutations of SARS CoV-2 result in decrease in the efficacy of the vaccines that have been developed before the emergence of these mutations. Thus, it is believed that using additional measures to combat the virus is not only advisable but also beneficial. Two antiviral drugs were authorized for emergency use by the FDA, namely Pfizer’s two-drug regimen sold under the brand name Paxlovid, and Merck’s drug Lagevrio. Pfizer’s two-drug combination consists of nirmatrelvir, a protease inhibitor that blocks coronavirus ability to multiply and another antiviral, ritonavir, that lowers the rate of drug clearance to boost the longevity and activity of the protease inhibitor. Merck’s drug Lagevrio (molnupiravir) is a nucleoside analogue with a mechanism of action that aims to introduce errors into the genetic code of the virus. We believe the armament against the virus can be augmented by the addition of another class of enzyme inhibitors that are required for viral survival and its ability to replicate. Enzymes like nsp14 and nsp10/16 methyltransferases represent another class of drug targets since they are required for viral RNA translation and evading the host immune system. In this communication, we have successfully verified that the Methyltransferase Glo, which is universal and homogeneous methyltransferase assay can be used to screen for inhibitors of the two pivotal enzymes nsp14 and nsp16 of SARS CoV-2. Furthermore, we have carried out extensive studies on those enzymes using different RNA substrates and tested their activity using various inhibitors and verified the utility of this assay for use in drug screening programs. We anticipate our work will be pursued further to screen for large libraries to discover new and selective inhibitors for the viral enzymes particularly that these enzymes are structurally different from their mammalian counterparts.

## Introduction

The recent pandemic outbreak of COVID19 caused by the Severe Acute Respiratory Syndrome Coronavirus 2 (SARS-Cov-2) was first reported in Wuhan, China in late 2019 resulting in infection of about 550 million and death of over 6.3 million world-wide and over 87 million infections with over one million deaths in the USA as of July 2022 (WHO). The infection by this virus is highly transmissible and can lead to severe pneumonia and death. The SARS-CoV-2 is an enveloped virus and belongs to the beta coronavirus family containing a large and complex positive-sense single stranded RNA genome (~30kb) which is similar to the other known six human coronaviruses (SARS-CoV, MER-CoV, HKU1, NL63, OC43, and 229E). The virus binds to human ACE2 receptor with high affinity, compared to the other CoVs, via its spike protein (S) followed by membrane fusion leading to the release of its genomic RNA that replicates and produce higher number of subgenomic RNAs. The viral RNA is then capped to limit its degradation and enhance its translation in the host cell (1–4).

Because of excessive number of casualties of this virus in the USA and worldwide, which is magnified by the impact of the viral infection and its effect on the worldwide economies, there is tremendous interest in prevention of viral infection by minimizing individual contacts and following good hygiene practice till development of vaccines, or therapeutics that can neutralize the virus or to inhibit its replication. There have been several attempts worldwide by many biotechnology and pharmaceutical companies as well as academic and government institutions to develop vaccines for neutralizing the virus which resulted in the launch of very effective vaccines using mRNA of the S protein (Moderna and Pfizer/BioNTech) or using full inactivated virus using adenovirus carrier (Johnson and Johnson and AstraZeneca) (5). However, due to the increased mutating capability of the virus, new variants have emerged in many parts of the world casting doubts on the efficacy of these vaccines. These continuous emergence of variants of concern raise tremendous questions on the long-term efficacy of these vaccines on current and future mutating variants (6,7).

Several attempts were made to develop small molecule therapeutics that target the replicating machinery of the virus by developing inhibitors towards RNA-dependent RNA polymerase, proteases, and nucleases and more recently methyltransferases. Studies on repurposing drugs that have been used to treat HIV, HCV and other viruses to combat COVID19 was published recently by interim WHO Solidarity trial. These included remdesivir, lopinavir, interferon, and hydroxychloroquine. Unfortunately, even though remdesivir has been approved by the Food and Drug Administration (FDA) for minimizing the duration of hospital stay (8), and dexamethasone for reducing mortality in intensive care unit (ICU) patients by targeting inflammation (9), all have shown very little effect and currently there are no drugs against COVID-19 with efficacy proven in clinical trials. Most recently, Pfizer and Merck pharmaceuticals gained FDA approval for their therapeutic drugs targeting the SARS-CoV 2 proteases that mitigate the viral impact on patient health and minimize their hospitalization (10).

The SARS-CoV-2 genomic RNA is polycistronic with two large ORFs (1a and 1b) that encode several non-structural proteins (nsp) that include replicating enzymes, proteases, nucleases, and methyltransferases. In addition, the genome contains other structural proteins and accessory proteins. These proteins are generated using Cap-dependent translation and frameshifts (11). The viral RNA is targeted by the host immune system for which the virus acquired defense by mimicking canonical capping mechanism to cap the 5’ ends of the RNA molecules similar in function to the host capping enzymes, thus escaping the innate cellular immune response and replicate efficiently (11–13). In the case of coronaviruses, there are two main enzymes that methylate viral mRNA, these include nsp14 and nsp16. The enzyme nsp14 is a bifunctional enzyme with an exoribonuclease (ExoN) at its N-terminus for proofreading, and a Guanine-N7 methyltransferase activity (N7-MTase) at its C-terminus for capping viral mRNA. The other enzyme is nsp16 that methylates nucleoside-2’O (2’O-MTase) which in combination with nsp10 as a heterodimeric complex methylate 2’-ribose of the 5’ penultimate nucleotide. The initial Cap core structure (cap-0) is formed in multi steps resulting in the formation of Gppp-RNA which is methylated at the N7-position of the capped guanine by N-7-MTase and then by 2’-O methylation of the first nucleotide ribose resulting in the formation of cap-0 and cap-1 respectively. Thus, the 5’cap consists of 7-methylgunaosine (m7G) linked to a 5’-5’ triphosphate bridge followed by a methylated nucleotide 2’ hydroxyl group culminating in the formation of ^m7^GpppN_m2’o_ where N represents the first nucleotide after the capped G nucleotide.

Thus, the function of methylating 5’-end cap is to provide stability for mRNA and allow for ribosomal recognition by eucaryotic translational machinery that are critical for viral genome replication within a host cell cytoplasm (14,15). Furthermore, nsp16 and its activator nsp10 appear to be very promising molecular target from the perspectives of drug design because it is indispensable for replication of the virus in cell culture. While the homolog of nsp16 in humans (CMTr1) does the same reaction as nsp16, CMTr1 does not require another nsp protein (nsp10) for enhanced activity (16,17). Using molecular dynamics simulations, it was possible to show that nsp10 binding to nsp16 shifts its conformation to become active. Thus, nsp16 requires nsp10 as a stimulatory factor to carry out 2’-O-MTase activity (16–18). These structural differences between viral and host MTases can be exploited to develop antiviral drugs.

To facilitate the development of novel inhibitors against this virus, we took advantage of a biochemical Methyltransferase assay that we previously developed to monitor the activity of these two SARS-CoV-2 methyltransferases nsp10/16 and nsp14 and carry out validation studies using methyltransferase inhibitors including small molecules as well as peptides (19). Others have developed enzymatic assays (20–23) and/or performed virtual screening (24) to monitor methyltransferase activities in an attempt to identify inhibitors from chemical libraries (24). The assay presented here is bioluminescent, homogeneous, universal for all MTases, and HTS compatible to enable the search for novel inhibitors of these enzymes for future development into therapeutics (19).

## MATERIALS AND METHODS

### Cloning and Expression of Methyltransferases

Cloning and Expression of SARS-CoV-2 nsp 10, nsp 16, and nsp14 Methyltransferases was carried out using sequences available at Genbank MN908947.3 data bank and the genes were cloned and expressed in E. coli according to the protocol described by SignalChem Lifesciences Corp., Richmond, BC, Canada. Briefly, SARS-CoV-2 full-length nsp10, nsp14 and nsp16 (Genbank: MN908947.3) genes were respectively inserted into a pET-21b vector possessing a C-terminal 6X His tag. The plasmids were transformed into E. coli BL21(DE3). The transformed cells were cultured at 37°C in 20 mL LB media for overnight (14-16 hours). The starter cultures were then inoculated into 1-liter LB medium and grew at 37°C until the OD_600_ reaches 0.6 followed by Induction of protein expression by adding IPTG to a final concentration of 0.2 mM and incubated at 25°C for 4 hrs. The cells were harvested through centrifugation at 5000 rpm for 5 minutes and cell pellets were resuspended in lysis buffer (50 mM sodium phosphate, pH7.5, 300 mM NaCl, 1 mM β-glycerophosphate, 0.5% NP-40, 1mg/mL lysozyme, 10 μg/mL STI, 0.2 mM PMSF, 0.1 μM sodium vanadate, 0.1% β-mercaptoethanol, 10% glycerol) and homogenized by stirring at 4 °C for 30 minutes followed by 3 pulses of sonication (15 seconds each) on ice. The cell lysates were clarified through centrifugation at 10,000 rpm for 10 minutes. The proteins were purified using TALON Metal Affinity Resin (Takara Bio) and eluted with a buffer containing 200 mM imidazole, 50 mM sodium phosphate (pH7.5) and 300 mM NaCl. Pooled enzymes were dialyzed in 50mM Sodium Phosphate, pH 7.0, 300mM NaCl, 150mM Imidazole 0.1mM PMSF, 0.25mM DTT, and 25% glycerol and stored at −70°C.

### Chemicals and assay components

ATA: Aurintricarboxylic Acid, Sinefungin, and Dimethyl sulfoxide (DMSO) were purchased from Millipore-Sigma Corp. (St. Louis, MO). Peptide P29 (YGG ASV CIY CRS RVE HPD VDG LCK LRG KF-NH2) was synthesized by AnaSpec Inc./Kaneka Eurogentec (EGT Corporate, Fremont, CA). Methyltransferase reaction buffer: 20mM Tris buffer, pH 8.0, 50mM NaCl, 1mM EDTA, 3mM MgCl2, 0.1mg/ml BSA, and 1mM DTT.

Methyltransferase-Glo™ Assay kit from Promega Corporation (Madison, WI) contains Methyltransferase-Glo™ Reagent, Methyltransferase-Glo™ Detection Solution, SAH (S-Adenosyl-Homocysteine), and SAM (S-Adenosyl-Methionine).

Assay plates: Commercial assay plate for 96-well and low volume 384-well: Assay plate 96-well and low volume 384-well non-Treated white polystyrene or EIA/RIA plate 96 well Half Area non-Treated white polystyrene (Corning Costar: 3912, 3693, 3572, and 4512, Corning Incorporated, Corning, NY), or equivalent.

### Oligonucleotides Substrates

The RNAs 1 to 4 were purchased from New England Biolab (Ipswich, MA, USA): RNA#1: G(5’) ppp(5’) A RNA Cap Structure Analog (S1406); RNA #2: m7 G(5’) ppp(5’) A RNA Cap Structure Analog (S1405); RNA #3: G(5’) ppp(5’) G RNA Cap Structure Analog (S1407); and RNA #4: m7 G(5’) ppp(5’) G RNA Cap Structure Analog (S1405). Other oligonucleotide (oligo RNAs) substrates were synthesized by TriLink Biotechnologies (Maravai LifeSciences, San Diego, CA, USA): Oligo #1: 5’ {Gppp.}A[mG] UUG UUA GUC UA 3’; Oligo #2: 5’ {Gppp.}AG UUG UUA GUC UA 3’; Oligo#3: 5’ {N7-MeGppp.}AG UUG UUA GUC UA 3’. Additional Oligo RNAs were obtained from IDT Technologies, Inc (Coralville, Iowa, USA): Oligo RNA 0011: 5’-UAC ACU CGA UCU GGA CUA AAG CUG CUC-3’; Oligo RNA 0012: 5’-ACG AGU CCU GGA CUG AAA CGG ACU UGU-3’; Oligo RNA 0020: 5’-ACA GAG A-3’; and Oligo RNA 0021: 5’-UAC AGA GAA-3.

### SAH Standard Curve

SAH standard curve was prepared in the Methyltransferase reaction buffer to assess the linearity of the assay during enzyme titrations in order to calculate the amount of SAH produced at each concentration of enzyme used. The 1μM and 10μM SAH concentration stock were made first in Methyltransferase reaction buffer. To perform ½ dilution factor of SAH, 200μl of SAH solution (1μM or 10μM) was added to well A1 and 100μl of Methyltransferase buffer was added into well A2 to well A12 in a regular 96-well plate. A serial twofold dilution was performed by transferring 100μl from well A1 to well A2 and mix by pipetting up and down. Then, 100μl was transferred from well A2 to well A3 and the transfer/mix process was repeated from well A3 through well A11. SAH solution was then discarded and well A12 contained only buffer. Once standards are prepared, 4μl of each of the 12 points was transferred to the assay plate and SAH was detected using the Methyltransferase-Glo™ Assay following the manufacturer’s procedure. Briefly, 1μl of 0.5X Methyltransferase-Glo™ Reagent was added then the plate was mixed for 1-2 min and incubated at room temperature (23° C) for 30 min. Then, 5μl Methyltransferase-Glo™ Detection Solution was added and the plate was mixed again before incubating it for another 30 min to be used by luciferase/luciferin reaction. Luminescence was recorded on a plate reading luminometer.

### Methyltransferase Assay Conditions

Generally, all methyltransferase reactions were performed in methyltransferase reaction buffer using 384 low volume solid white assay plates described previously (18).

For SARS-CoV-2 methyltransferase reactions were performed at 37°C in 4μl for 90min with a proper plate seal. However, for the kinetic studies to determine substrate Km values, the reactions were performed at room temperature (23°C) in 4μl, followed by addition of 1μl of TFA to a final concentration of 1% to stop the reactions. Then, 5μl Methyltransferase-Glo™ Detection Solution was added and the plate was mixed again before incubating it for another 30 min to be used by luciferase/luciferin reaction and Luminescence is recorded.

### Inhibitor Studies Using Methyltransferase-Glo™ Assay

For determination of IC_50_ of inhibitors for SARS-CoV-2 methyltransferases, reactions were performed in 5μl mixture containing 1X methyltransferase reaction buffer, with or without 0.4% DMSO, substrate RNA and either 1ng nsp14 or 60ng nsp10/nsp16 per reaction containing 5μM, 25μM, or 50μM of SAM, and serial dilution of small molecules to be tested at each SAM concentration. Reactions were incubated at 37°C for 90 minutes. After incubation time, MTase-Glo assay protocol was applied before luminescence was recorded and IC_50_ values were determined. To test P29 polypeptide for inhibiting methyltransferase activity, reactions containing 1ng of nsp14 were carried out same as previously mentioned for small molecule inhibitors. However, with nsp 10 and nsp16 study, the nsp10 and nsp 16 were tested individually or in combination. Inhibition of nsp16 by P29 was carried out by pre-incubating P29 with or without nsp16 for 10min to 15min at room temperature (23°C) before mixing reactions with nsp10 at the following nsp16:nsp10 ratios: 0μM (1:0), 0.5μM (1 : 1), 1.0μM (1 : 2), 2μM (1 : 4), and 3μM (1 : 6) and reactions were started by adding substrates (oligo RNAs and SAM) with MTase-Glo Reagent. The reactions were incubated at 37°C for one and half hour. before SAH was detected using MTase-Glo assay protocol.

### Signal Detection and Data Analysis

All low volume 384-well and regular 384-well assay plated were read using infinite 500 or M1000PRO instruments from Tecan Ltd. (Mannedorf, Switzerland). For plotting, analysis of data and calculation of methyltransferases biochemical parameters, both Microsoft Excel and Prism from GraphPad Software (La Jolla, CA) were used. IC_50_ values were determined by using a nonlinear regression fit to a sigmoidal dose response (variable slope).

## Results and Discussion

To demonstrate that the assay we developed can determine the activity of SARS-CoV-2 methyltransferases (nsp14 and nsp10/16) and is amenable to high throughput screening for potential methyltransferase inhibitors that may progress forward to clinical testing, it was necessary to establish the sensitivity and linearity of the assay performance.

As we have already demonstrated, the assay principle is based on the detection of S-adenosyl homocysteine (SAH), a universal product of all methyltransferases using S-adenosylmethionine (SAM) as the universal substrate. The assay principle is depicted in Figure 1A where the substrate SAM is utilized by the methyltransferase and generating SAH which is converted in sequential reactions to ATP which can be quantitated using luciferase/luciferin bioluminescence reaction. The amount of light generated (RLU) is proportional to the SAH concentration. As shown in Fig 1B-C, the assay can detect as low as 30 nM of SAH and is linear up to 10 μM of generated SAH. Thus, we verified that the assay is not only specific since the control without SAH showed very low luminescence, but this linearity and high sensitivity will allow the detection of methyltransferases with broad range of specific activities.

**Figure 1.**
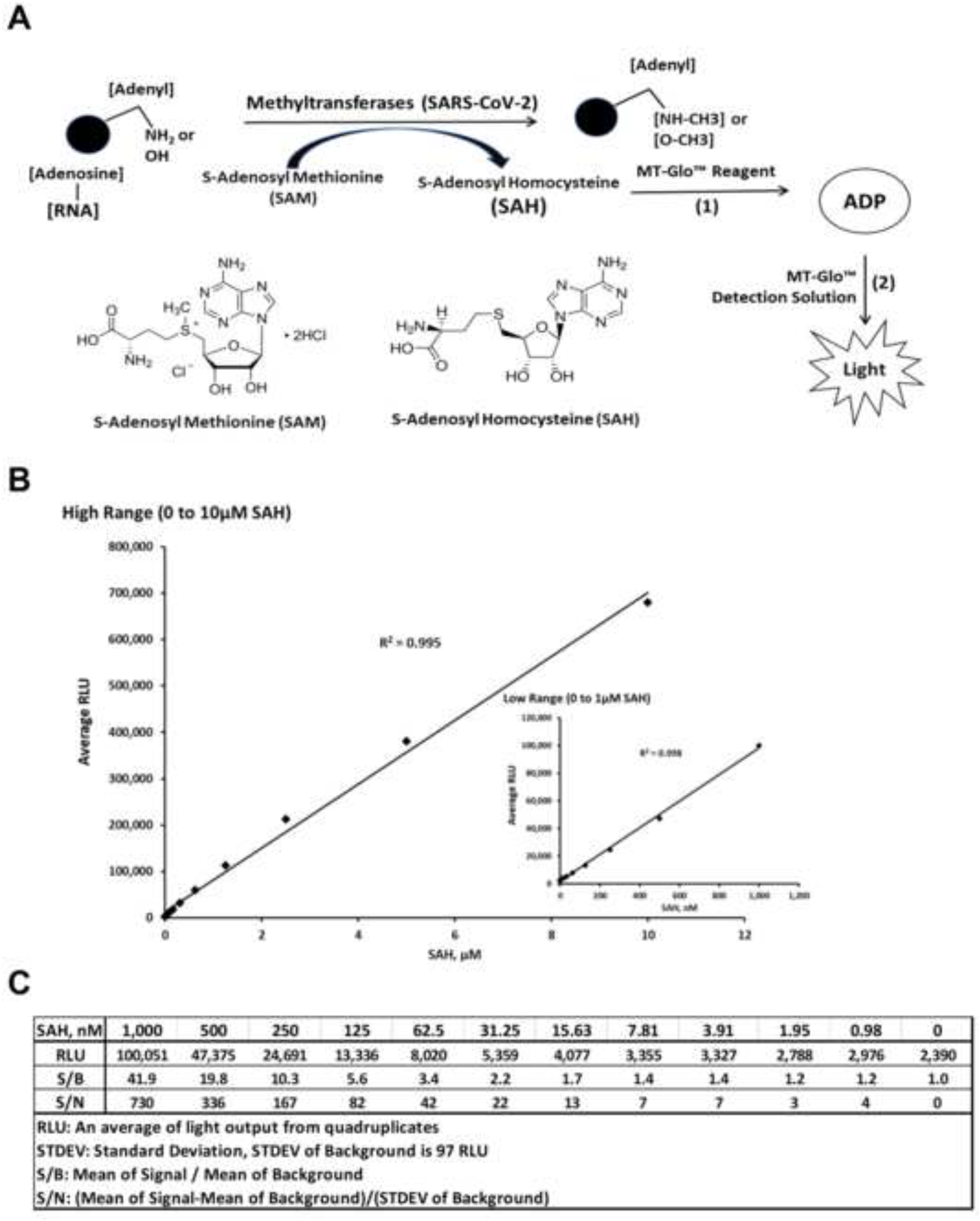
Universal MTase-Glo™ Scheme and Sensitivity of SAH Detection. (A) Schematic diagram of the basic principle of MTase Glo^TM^. Basically, any methyltransferase (MTase) that is capable of methylating its substrate and S-Adenosylmethionine (SAM) will generate methylated substrate and S-Adenosylhomocysteine (SAH), the latter is a universal product for all methyltransferases. Conversion of SAH to ADP and to ATP is performed by coupled enzyme reactions. The ATP produced can be quantitated by a luciferase/luciferin bioluminescent reaction. (B) A standard curve showing the relative luminescence units (RLU) in response to SAH concentrations, inset show a lower range of SAH concentration vs. RLUs. Reactions were assembled with the indicated concentrations of SAH in a low-volume 384-well white solid plate. The MTase-Glo™ Assay was performed as described in Materials and Methods. Data were collected using a plate reading luminometer (INFITIE M1000PRO, TECAN). Each point represents average of four data points; the error bars represent the standard deviation. Data analysis was performed with Excel, Microsoft Office 365. (C) Table representing the data generated to quantify SAH concentrations and calculating signal to background and signal to noise ratio from panel B.

### Testing SARS-CoV2 Methyltransferases for selectivity towards RNA substrates: Substrate Selectivity for nsp14

Since the enzyme nsp14 is known as a capping RNA methyltransferase by methylating the 5’Guanisine triphosphate, we checked its activity using multiple capped and uncapped RNA substrates, in order to find out whether the presence of methylated capped RNA will be as a good substrate as the unmethylated uncapped RNA. For this, we titrated nsp14 in the presence of two groups of substrates. The first group consists of only dinucleotides and the second group contained longer oligonucleotide that are either uncapped or methylated and capped).

As shown in Figure 2A, in the presence of **50** μM SAM and 40 μM Oligo RNAs, SARS-CoV-2 nsp14 methyltransferase generated the highest activity with the uncapped RNA #1 substrate (G (5’) ppp(5’) A RNA) with as little as 5 ng of nsp14 producing a maximum activity. However, when we used the analog substrate RNA# 2 (m7 G(5’) ppp(5’) A) which is identical to RNA#1 but capped, it was a poor substrate with as high as 80 nanogram of nsp14 generating only one sixth of its activity compared to nonmethylated RNA#1 substrate. This suggests that nsp14 prefers uncapped substrates which is expected since its main function is to methylate the 5’G nucleotide. To test whether the nature of the second nucleotide following the first G nucleotide has any effect on the enzyme preference for the substrate, we titrated nsp14 as described above but using the RNA substrate G (5’) ppp(5’) G (RNA#3) instead of G(5’)ppp(5’)A and as a control, its capped version m7 G(5’) ppp(5’) G (RNA#4). It is apparent that the enzyme methylated the oligonucleotide when G in the penultimate nucleotide with similar high activity as when A was in the penultimate position. This means that the nsp14 enzyme loses its stringent selectivity of methylating only the noncapped oligo when the penultimate nucleotide was G albeit at lower rate than with the other noncapped oligonucleotide shown in Figure 2A. In other words, nsp14 can have activity towards methylated RNA if the penultimate nucleotide is G.

**Figure 2.**
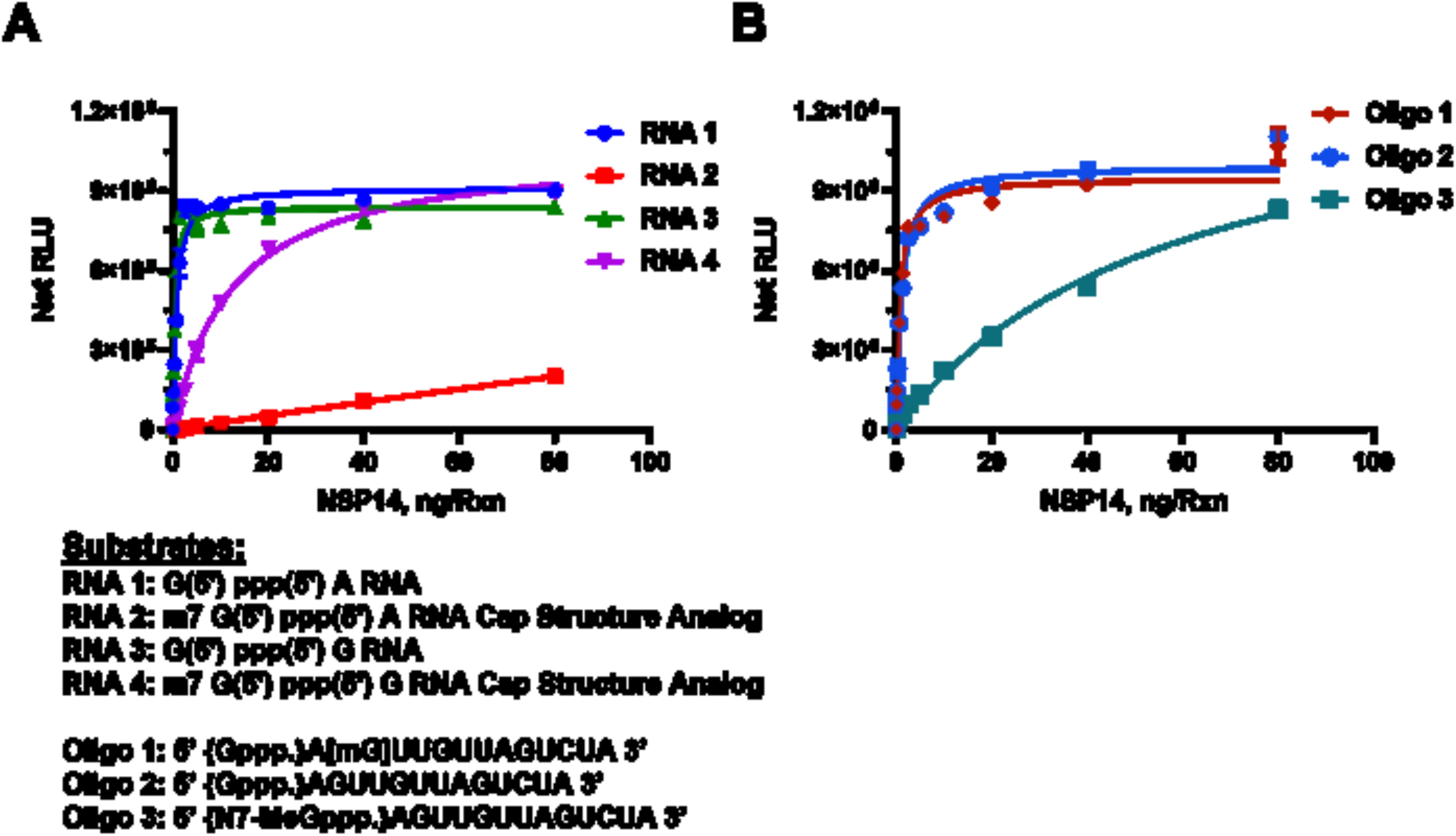
Determining SARS-CoV-2 nsp14 Methyltransferase Activity using RNA Substrates. Monitoring the activity of nsp14 methyltransferase using capped and uncapped RNA substrates as listed under the graphs. (A) Short sequences substrates, and (B) long sequence substrates. Reactions were carried out with 50μM SAM and 40μM of either short RNAs or Oligo RNAs using a white solid 384LV-well plate. SARS-CoV-2 nsp14 reaction was for 90 min at 37°C. The list of RNA substrates and their sequence are shown below the figures. The MTase-Glo™ Assay was performed as described in Materials and Methods section. Each point represents an average of two data points; the error bars represent the standard deviation. Data analysis was performed with GraphPad Prism^®^ software, version 9.1.0, for Windows^®^ using a One Site Binding (hyperbola) program.

To understand fully the substrate requirement for the enzyme, we tested its activity using longer oligonucleotides. The enzyme was again titrated in the presence of an uncapped substrate, a methylated substrate on the 5’G nucleotide, or methylated at the second position nucleotide. As shown in Figure 2B, the enzyme was active against the nonmethylated substrate as well as the substrate methylated nucleotide at the 2’of the penultimate position. As expected, its activity was much lower when the substrate used was capped (Oligo#3). This suggests that as long as the 5’nucleotide is not methylated, the oligo serves as a good substrate even if the methylation is on the second position on the oligo substrate. It is noteworthy that the enzyme has similar activity using either shorter oligo (dinucleotides) (Fig 2A) or longer substrates (Fig 2B). It is also evident that the enzyme shows low activity when the substrate is methylated at the 5’ nucleotide. It appears that the nsp14 enzyme shows 100-fold less activity with methylated capped than nonmethylated uncapped substrates.

To assess the affinity of nsp14 towards its substrates, we measured the nsp14 methyl donor and acceptor substrate apparent kd values. We compared the activity of the enzyme towards short and long oligonucleotide substrates using different concentrations of both substrates in the presence of constant concentration of the enzyme and the methyl donor SAM (50μM) at 23°C for 30min. It appears that the enzyme prefers the short sequence since apparent kd for the substrate is 1.01 μM and 1.74 μM for the short vs the long oligonucleotides, respectively (Figure 3A).

**Figure 3.**
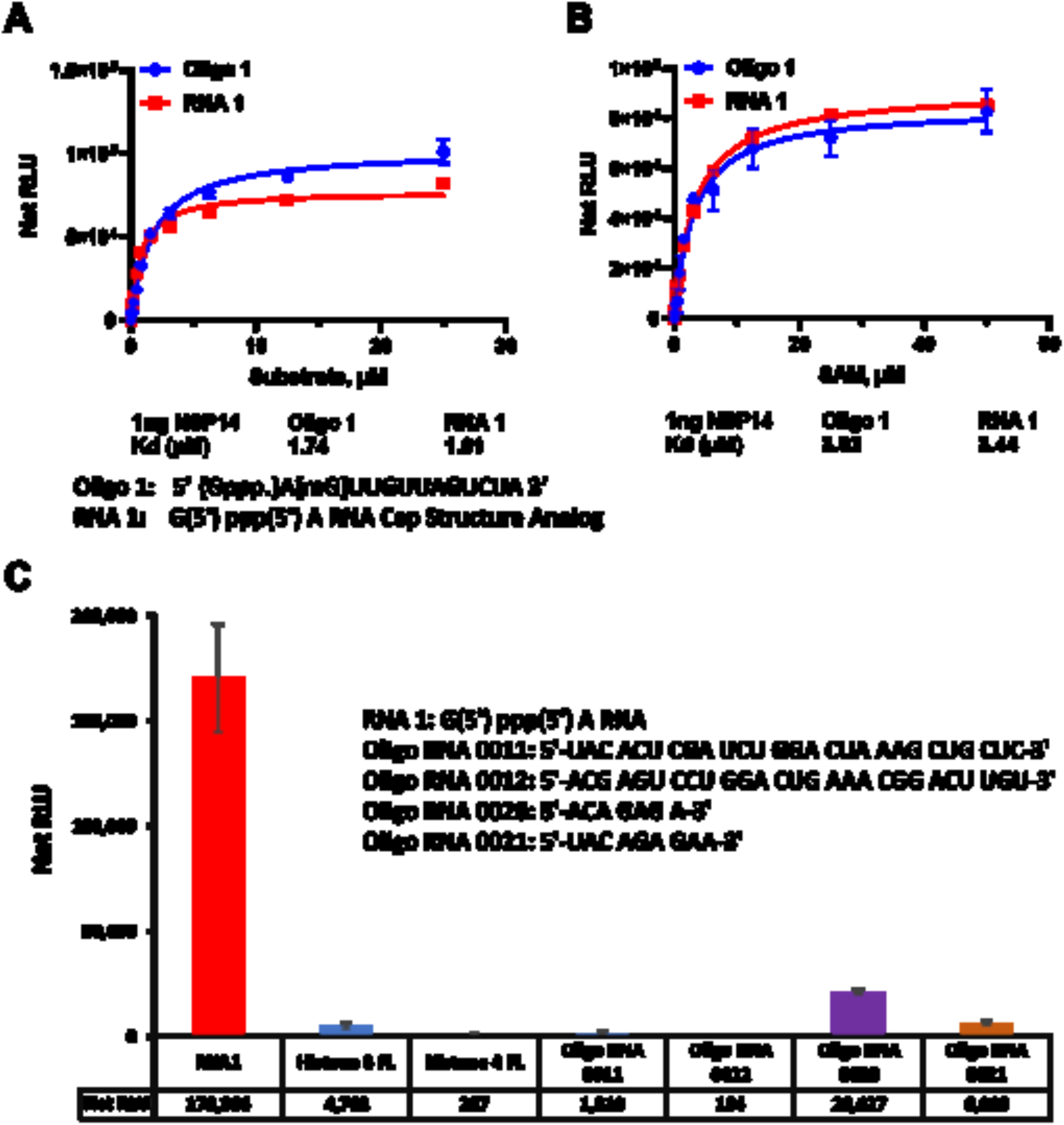
SARS-CoV-2 nsp14 Methyltransferase Kinetic Parameters and Substrate Specificity. (A) SARS-CoV-2 nsp14 enzyme activity was monitored using a titration of uncapped (RNA1) short sequence and uncapped oligo 1 but methylated at the penultimate nucleotide sugar in the presence of 50μM SAM per reaction. (B) Titration of SAM substrate using saturating concentrations of RNA substrates, 25 μM of oligo 1 and 50 μM of RNA1. The data were generated using 1ng of nsp14 per reaction at 23°C for 30min. (C) Substrate selectivity of nsp14 using different RNA sequences and proteins known to be methylated. Reactions were carried out using 1ng of nsp14 per reaction with 50μM SAM and 25μM of each Oligo RNA and 20μM for both Histones and reactions were carried out at 37°C for 80min in white solid 384LV-well plate. MTase-Glo™ Assay was performed as described in Materials and Methods section. Each point represents an average of two data points; the error bars represent the standard deviation. Data analysis was performed with GraphPad Prism^®^ software, version 9.1.0, for Windows^®^ using a One Site Binding (hyperbola) program (panel A and B) and with Microsoft^®^ Excel 365 program (panel C). All substrate sequences are shown in their specific panel.

We further tested the enzyme for its affinity towards the other substrate (SAM). As shown in Figure 3B, the enzyme has similar affinity for SAM in the presence of either substrate with apparent kd value around 3.9 μM. Thus, both substrates are methylated equally well with similar affinity for the enzyme towards SAM substrate. Thus, we conclude from these studies that the enzyme prefers shorter RNA Oligonucleotides and can tolerate the G at the penultimate position instead of A and can also methylate oligonucleotides that are methylated on the 2’ on the penultimate nucleotide.

Finally, because this methyltransferase assay can be used with a wide range of substrates and substrate concentrations, we compared the enzyme selectivity towards RNA oligonucleotides as well as oligopeptides that are known to be substrates for other methyltransferases. We used a full-length histone H3 and full-length histone H4 that are known substrates for protein methyltransferases. As shown in Figure 3C, it is evident that the enzyme showed no activity towards protein substrates. We also tested two oligonucleotides known to be substrates for adenine N-6 methyltransferases MTTL3/14 (Oligo 0011 and 0012) and we could observe only low activity towards those substrates. Similarly, other substrates for RNA methyltransferase MTTL 16 (oligo 0020 and 0021) showed some activity with nsp14. The substrate selectivity supports the notion that nsp14 does not utilize substrates of the MTTL RNA methyltransferase class of enzymes.

### Substrate selectivity for SARS-CoV-2 nsp10/16

Since SARS-CoV-2 methyltransferase nsp10/16 methyltransferase heterodimer is known to methylate the ribose sugar of the penultimate nucleotide of RNA substrate, we used methylated capped oligonucleotide (Oligo#3) and other substrates to assess the enzyme activity. Similarly, to nsp14, we observed that the enzyme shows high selectivity towards small oligo and other oligonucleotides that we tested for nsp14 (Fig 4A) but not MTTL 3/14 substrates or histone protein substrates. However, nsp10/16 showed reasonable activity with MTTL16 substrates with almost half of its activity when using oligo 3 substrate.

Moreover, the results in Fig 4B show that the enzyme methylates the oligonucleotide #3 well with an apparent Kd of 10-11 μM. Similarly, the enzyme has high affinity for SAM as substrate like many other methyltransferases with apparent kd of 33 μM (Fig 4C).

**Figure 4.**
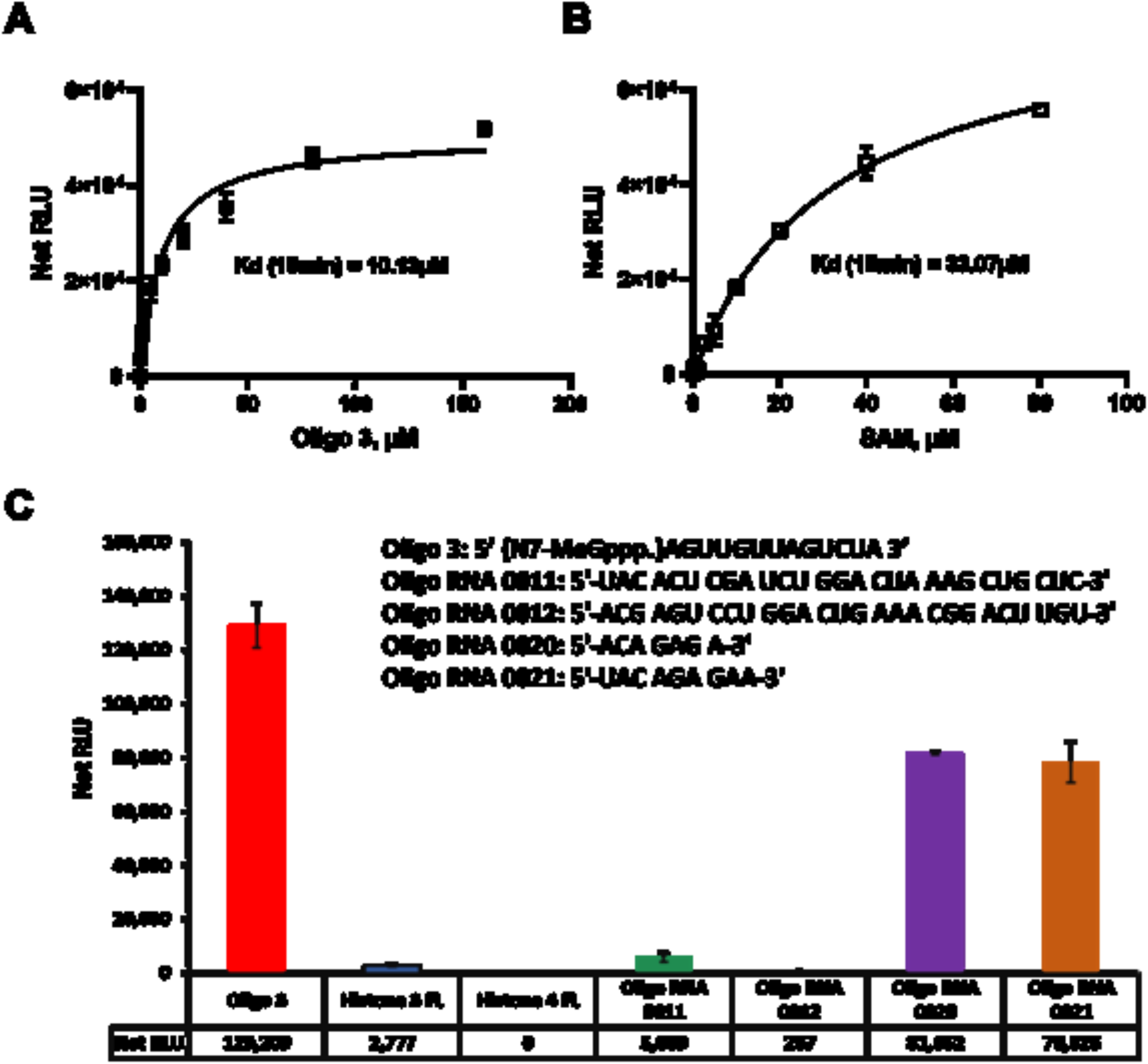
SARS-CoV-2 nsp10/nsp16 Methyltransferase Kinetic Parameters and Substrate Specificity. (A) SARS-CoV-2 nsp10/nsp16 substrate kinetic studies were carried out using different concentrations of Oligo 3 using 20μM SAM per reaction. (B) Similar studies with the enzyme using 80 μM of Oligo 3 and varying SAM concentration to calculate the Kd value for SAM. The kinetic studies were carried out with 40ng of nsp10/nsp16 complex per reaction at 23°C for 15min. (C) Testing substrate specificities for nsp10/nsp16 enzyme reactions were carried out using 40ng enzyme per reaction with 50μM SAM and 25μM of each Oligo RNAs and 20μM for both Histones. Reactions were carried out at 37°C for 80min in white and solid low-volume 384-well plate, and the MTase-Glo™ Assay was performed as described in Materials and Methods section. Each point represents average of two data points. Data analysis was performed with GraphPad Prism^®^ software, version 9.1.0, for Windows^®^ using a One Site Binding (hyperbola) program (panel A and B) and with Microsoft^®^ Excel 365 program (panel C). All substrate sequences are shown in their specific panel.Data analysis was performed with Microsoft^®^ Excel 2013 program, and GraphPad Prism^®^ software, version 9.1.0, for Windows^®^ using a One Site Binding (hyperbola) program

### Determination of the heterodimerization ratio of nsp10 and nsp16 on methylation of substrate

It has been documented that replication and translation of RNA viruses in eucaryotic cells requires capping of RNA by capping viral methyltransferase (cap-0) followed by methylation of capped RNA at the 2’-ribose of the penultimate nucleotide resulting in the formation of cap-1 structure that mimics cellular mRNA and thus preventing recognition of the viral RNA by host innate immunity (25–26). As mentioned earlier, with SARS-CoV-2, the first capping MTase is nsp-14 and the second MTase is nsp16 resulting in the formation of cap 0 and cap1 respectively. Nsp10 interacts exclusively with the ExoN domain of nsp14, as is consistent with previous biochemical results showing that nsp10 stimulates the ExoN activity without perturbing the N7-MTase activity (27–30). In contrast, mutants in which the nsp10-nsp16 interaction was disturbed proved to be crippling but viable. These experiments imply that the nsp10 interaction with nsp14 and nsp16 and possibly other subunits of the viral replication complex may be a target for the development of antiviral compounds against pathogenic coronaviruses. Because nsp10 is encoded in about three-to sixfold excess over nsp14 and nsp16, the nsp14–nsp10 complex and nsp16–nsp10 complex can exist simultaneously (27–29). It is noteworthy that nsp10 is found in all coronaviruses (notably absent among prokaryotes or eukaryotes), and its sequence is highly conserved across the entire length of the protein suggesting its importance in the coronavirus life cycle (28).

As discussed earlier, although the human homolog of nsp16 (CMTr1) does the same reaction as the, viral counterpart nsp16 does not require another nsp protein (nsp10) for enhanced activity. Using molecular dynamics simulations, it has been shown that nsp10 binding to nsp16 shifts its conformation to be active (15), and that nsp10 is required for nsp16 to bind both SAM and RNA substrate and stabilizing the SAM binding pocket and extending the RNA binding groove of nsp16. Thus, nsp16 requires nsp10 as a stimulatory factor to carry out 2’-O-MTase activity. It was also reported that the ratio of nsp10 to nsp16 to form the complex could vary from 1:1 to 8:1 (nsp10: nsp16) and the higher nsp10 ratio increases the activity of MTase activity of nsp16 (25–27). It is believed that the physiological ratio of nsp10 to nsp16 is around 6:1 (28–30). Based on this literature findings, we tested the activity of nsp16 in the absence and presence of different concentrations of nsp10. As shown in Fig 5A, the activity of nsp 16 was enhanced significantly by the addition of nsp10 with all the SAM and Oligo#3 concentrations tested. It is interesting that 1:1 ratio of nsp10: nsp16 gave maximal 6-fold MTase activity with no further activation observed with higher nsp10. It is also interesting the MTase activity of nsp16/nsp10 was maximal at 25 μM SAM and 50 μM oligonucleotide. What is very clear from the literature is that nsp10 plays a modulatory role via its interaction with nsp14 and nsp16 via surface interactions with both proteins, but it does not have intrinsic enzymatic activity on its own. However, testing nsp10 for MTase activity in the absence of nsp16, we observed methyltransferase activity albeit is not at the higher enzymatic activity we observed when combined with nsp16. What was intriguing is that nsp10 alone shows MTase activity and is proportional to the enzyme concentration with maximal activity at 5 μM SAM and 100 μM of oligonucleotide 3 (Fig 5B).

**Figure 5:**
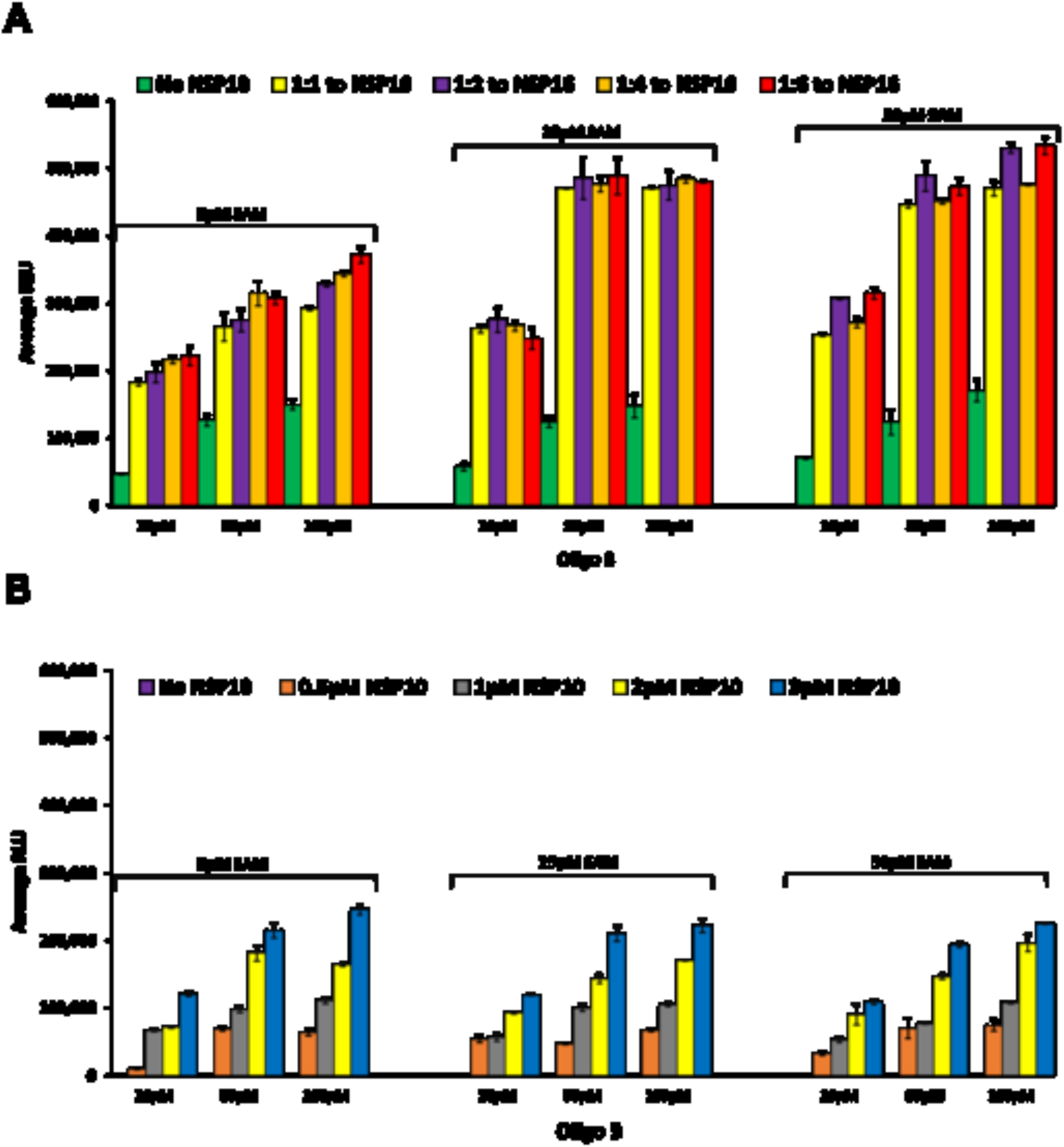
Determining Optimal Stoichiometry of SARS-COV-2 nsp10 & nsp16 Methyltransferases. (A) Monitoring methyltransferase activity of nsp10/nsp16 using constant concentration of nsp10 (0.5μM) with increasing concentration ratio of nsp16 (0, 1:1, 1:2, 1:4, and 1:6) in the presence of 5, 25, and 50μM SAM and different concentrations of Oligo 3 (10, 50, and 100μM). (B) Testing for methyltransferase activity of different concentrations of nsp10 only without nsp16. Enzyme activity was determined in the presence of different concentrations of SAM (5, 25, and 50μM) and different concentrations of Oligo 3 (10, 50, and 100μM). Reactions were carried out in white solid low volume 384-well plate for 90min at 37°C. MTase-Glo™ Assay was performed as described in Materials and Methods section. Each point represents an average of three data points; the error bars represent the standard deviation Data analysis was performed with Microsoft^®^ Excel 365 program.

### Inhibition of SARS-CoV2 methyltransferases with small molecules Effect of nsp10-derived peptide on nsp10/16 MTase activity

Since nsp10 is required for maximal activity of nsp16, it was suggested that a peptide derived from nsp10 can be used as inhibitor of nsp16 MTase activity (27–29). We synthesized P29 polypeptide that was reported to inhibit MTase activity of nsp16 and tested its potential inhibitory effect. We found that 200 μM of P29 inhibited the MTase activity of nsp16 (0.5μM) by 20% and 300 μM inhibited MTase activity by 40%. The percent inhibition was similar when using nsp10: nsp16 ratio of 4:1 and 6:1 (Fig 6A). It is noteworthy that percent inhibition was decreased by increasing oligonucleotide substrate concentration and loss of inhibition was observed when the substrate concentration increased to 100 μM suggesting a competition between the substrate and the nsp10 derived peptide for the enzyme nsp16 (Fig 6B).

**Figure 6.**
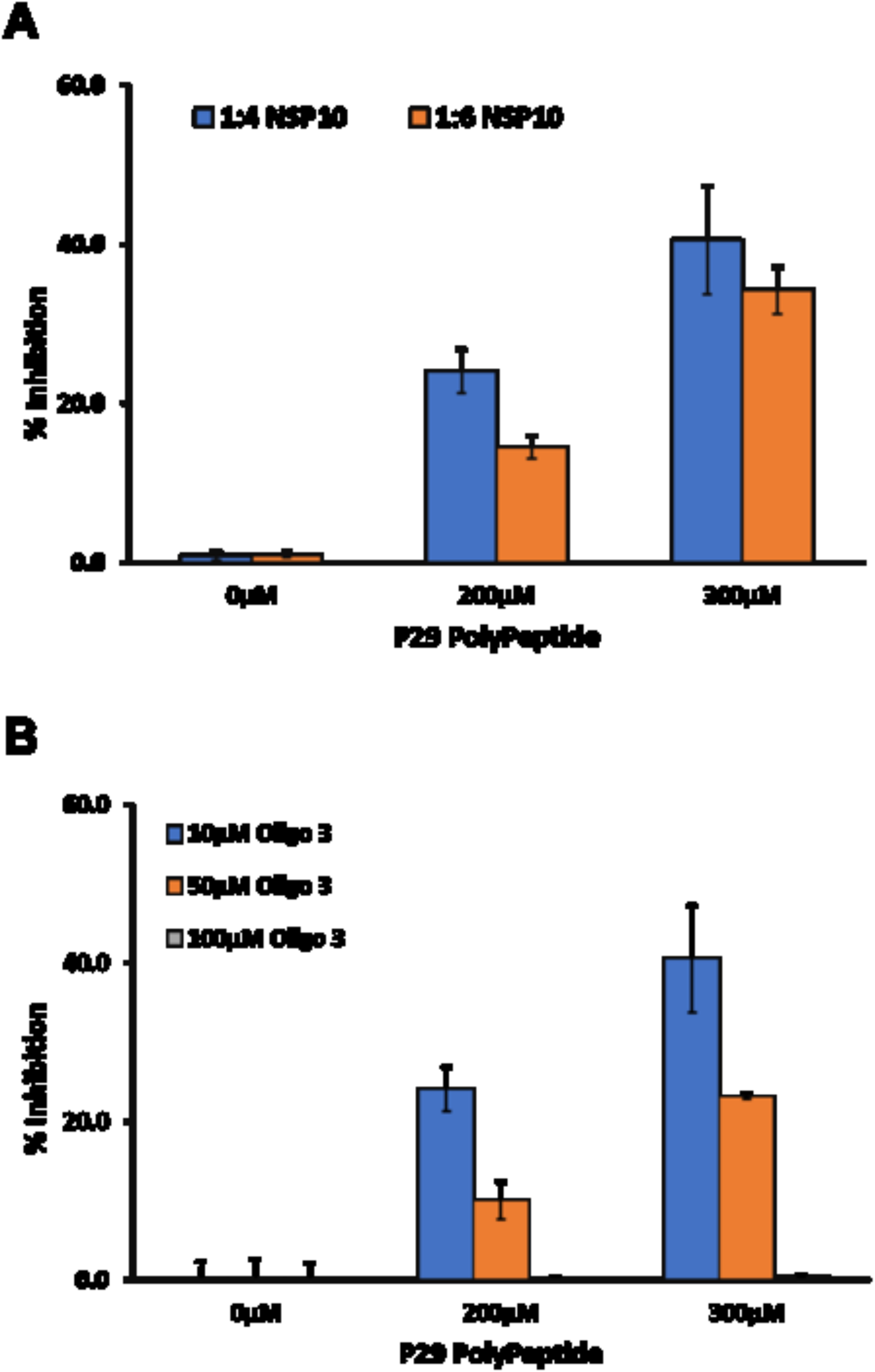
Effect of nsp10-derived peptide P29 on SARS-CoV-2 nsp10/nsp16 Methyltransferase activity. **Panel** A Inhibition studies of methyltransferase activity of (1:4, 1:6) of nsp16/nsp10 (0.5 μM:2.0 μM 0.5 μM:3.0 μM) using 0, 200 μM and 300 μM P29 on in the presence of 5μM SAM and 10μM Oligo 3. (B) Inhibition of nsp 16/nsp10 activity with varying concentrations of P29 using 0.5μM of nsp16/ with 2μM nsp10 (1:4 ratio) in the presence of 5μM SAM and increasing concentration of Oligo 3 (10, μM, 50 μM, and 100 μM). Reactions were carried out by preincubating nsp16 with P29, then started the reactions by the addition of nsp10 and incubated for 90min at 37°C. All reactions were performed in a white solid 384LV-well plate, and The MTase-Glo™ Assay was performed as described in Materials and Methods section. Each point represents an average of three data points; the error bars represent the standard deviation Data analysis was performed with Microsoft^®^ Excel 365 program.

### Effect of small molecule inhibitors on SARS-CoV-2 nsp14 MTase activity

It has been proposed that Aurintricarboxylic acid (ATA) and its ammonium salt shows antiviral activity in vitro against coronaviruses such as SARS, MERS and SARS-CoV-2, and thus proved useful in scientific research into novel antiviral drugs to combat these diseases. Similarly, the compound sinefungin has been frequently used as MTase inhibitor due to its competitive inhibition with SAM substrate (30–34). Therefore, we tested these two compounds against nsp14 using RNA1 and Oligo 1 substrates. Using RNA1 as a substrate, the results show that Sinefungin inhibited nsp14 MTase activity with IC_50_ of 2.41, 5.02, and 11.37 μM at 5, 25, and 50 μM SAM concentrations respectively, indicating that sinefungin inhibits nsp14 in a competitive manner towards SAM substrate. When, we tested the effect of sinefungin on MTase activity of nsp14 using the other oligonucleotide Oligo1 substrate to assess its potency (Fig 7A) against SAM concentrations, we observed the same trend, i.e., competitive inhibition against SAM but the IC_50_ were lower where it was 2.03, 4.21, and 5.2 μM at 5, 25, and 50 μM SAM concentrations respectively (Fig 7B). Thus, the data show that the assay reproduces the literature reported values by confirming sinefungin as a competitive inhibitor against SAM using several oligonucleotide substrates.

**Figure 7.**
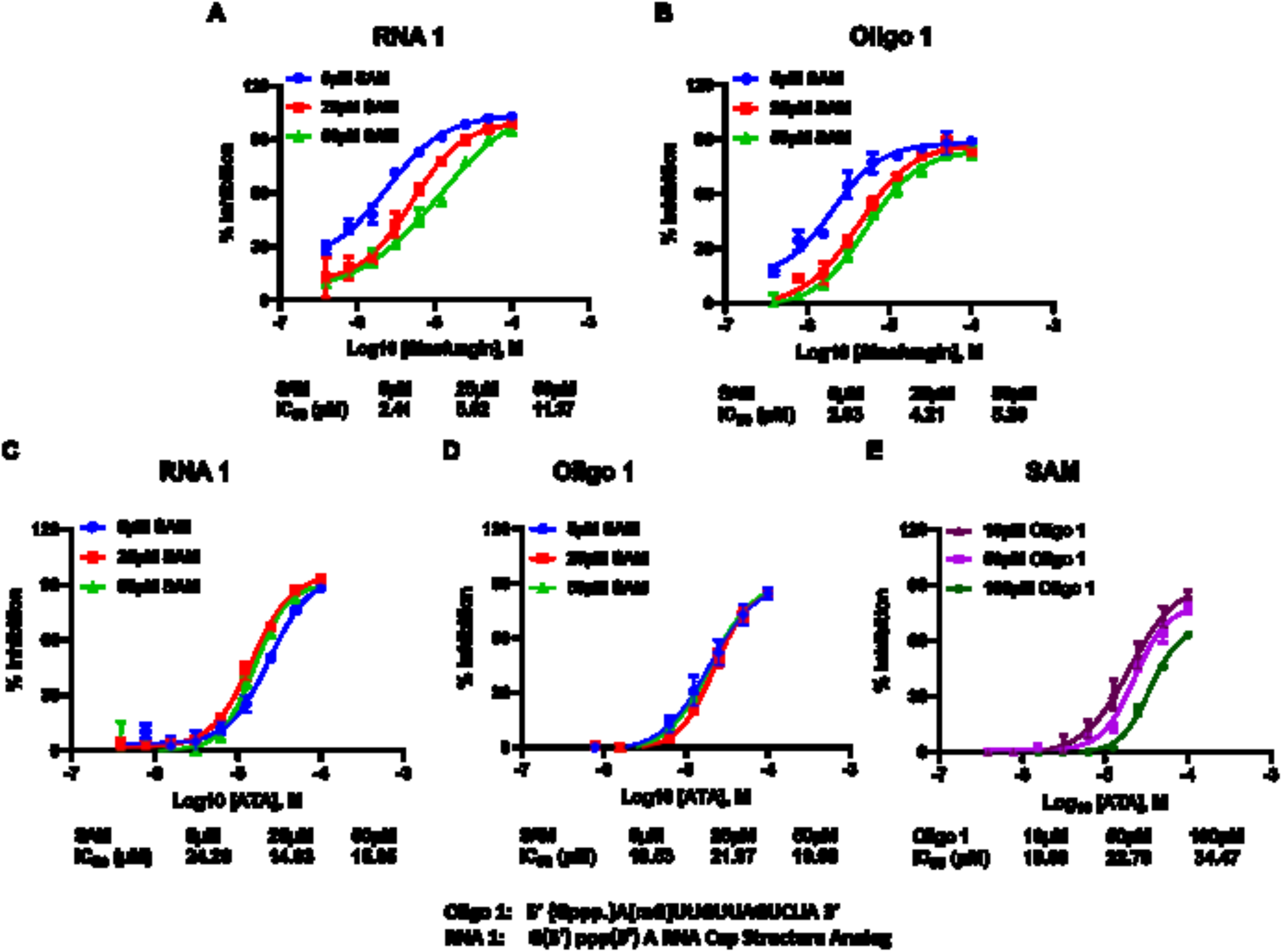
SARS-CoV-2 NSP14 Methyltransferase Inhibitors Study. Effect of sinefungin concentrations on nsp14 (1ng) methyltransferase activity at different concentrations of SAM (5 μM, 25 μM, and 50 μM) using 10 μM of RNA 1 (A) or 10 μM of Oligo1 (B). Effect of Aurintricarboxylic acid (ATA) on the activity of nsp14 (1ng) in the presence of varying concentrations of SAM (5 μM,25 μM, and 50 μM) using 10 μM RNA1 (C), or 10μM Oligo1 (D) and varying concentrations of Oligo1 (10 μM,50 μM,100 μM) at 5 μM SAM (E). Reactions were carried out for 90min at 37°C. All reactions were done using a white solid 384LV-well plate, and The MTase-Glo™ Assay was performed as described in Materials and Methods section. Each point represents an average of two data points; the error bars represent the standard deviation. Data analysis was performed with GraphPad Prism^®^ software, version 9.1.0, for Windows^®^ using a Sigmoidal dose response (variable slope) program.

While ATA was as potent inhibitor as sinefungin, it was not competitive with SAM since when tested against nsp14 using RNA1 and Oligo 1 substrates we observed no decrease in potency with increasing SAM concentration (Fig 7C-E). The IC_50_ of 24.36, 14.93, and 15.85 μM were obtained at 5, 25, and 50 μM SAM substrate respectively with RNA1 (Fig 7C). With the oligonucleotide oligo 1 substrate, ATA inhibited MTase activity of nsp 14 with IC_50_ of 19.53, 21.9, and 19.68 μM at 5, 25, and 50 μM SAM concentrations respectively (Fig 7D). This indicates its inhibition is not competitive with SAM concentrations in the reaction whether RNA1 or Oligo 1 substrate was used. However, when we tested ATA inhibition on nsp14 using different concentrations of Oligo 1, we see a slight effect of oligonucleotide substrate on the ATA inhibition of MTAse activity where IC_50_ increased from 19.6, to 22.70, and 34.47 μM at 10, 50, and 100 μM oligonucleotide substrate concentrations, respectively (Fig 7E). This may indicate that ATA could be competing slightly with the oligo substrate binding to nsp14.

### Studies on nsp10/16 MTases activity

We then tested the effect of these inhibitors on the MTase activity of nsp10/nsp16 (1:1). Using Oligonucleotide 3 and different concentrations of SAM, the activity was determined using 0, 50 μM, and 100 μM sinefungin. The results in Figure 8 show that sinefungin inhibited nsp10/16 in a similar manner to that against nsp14, showing competitive inhibition towards SAM. Using varying concentrations of Oligo 3 (0, 50μM, and 100μM) and 5 μM SAM, sinefungin has less inhibitory effect on MTase activity of nsp10/16. Thus, less competition by oligo 3 towards sinefungin than by SAM. Further studies are needed to explore the mechanism by which the peptide exerts its effect on MTase activity.

**Figure 8.**
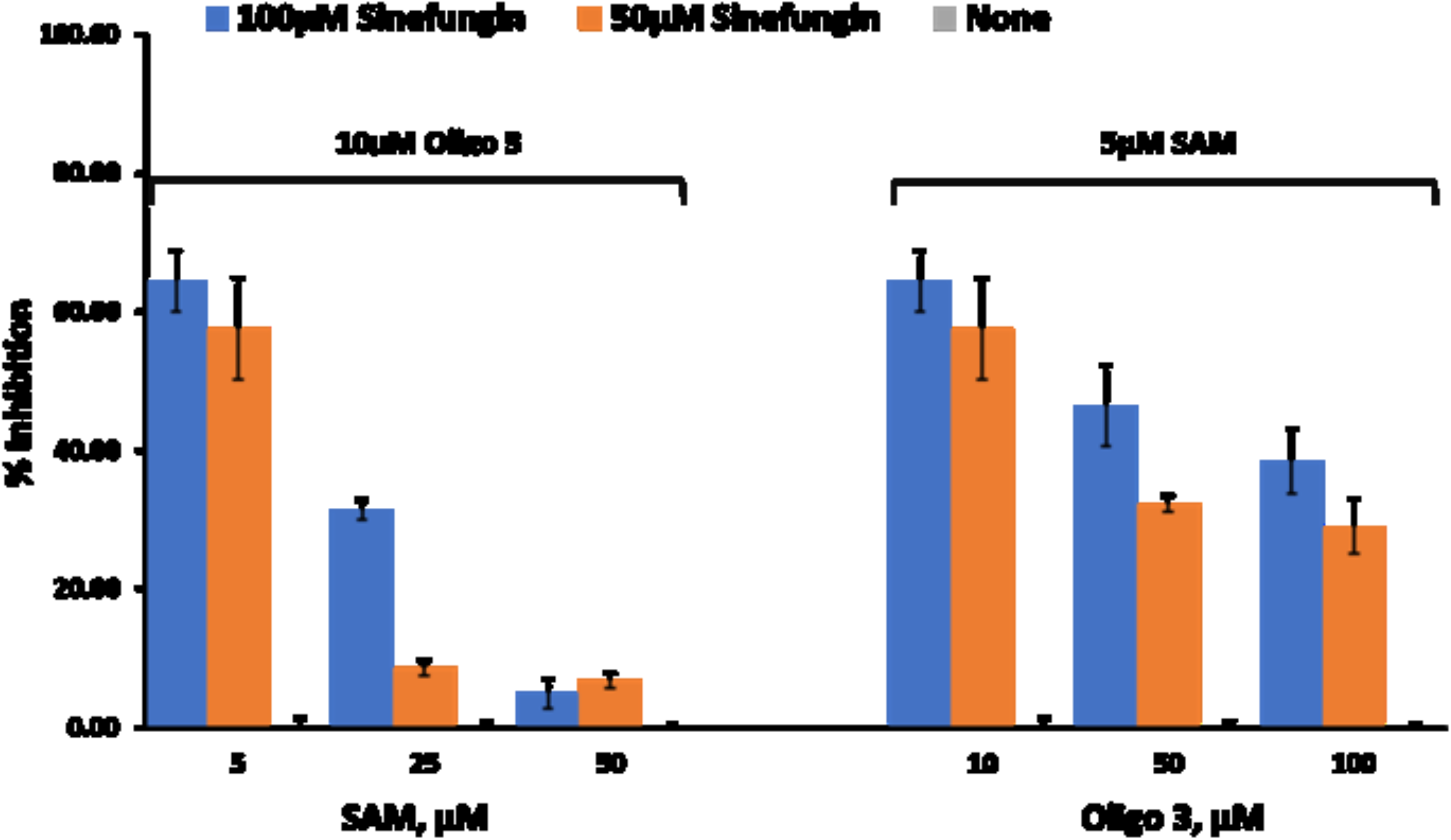
Effect of inhibitors on SARS-CoV-2 nsp10/nsp16 Methyltransferase activity. Percent inhibition of sinefungin (0 μM, 50 μM, and 100 μM) on the enzyme activity of SARS-CoV-2 nsp10/nsp16, at three different concentrations of SAM (5 μM, 25 μM, and 50 μM) using 10 μM oligo 3 as substrate (left panel) or different concentrations of Oligo 3 (10 μM, 50 μM, and 100 μM) using 5 μM of SAM. Reactions were carried out with 60ng of NSP10/NSP16 per reaction for 90min at 37°C. All reactions were done using white solid 384LV-well plate, and the MTase-Glo™ Assay was performed as described in Materials and Methods section. Each point represents an average of two data points; the error bars represent the standard deviation was performed with Microsoft^®^ Excel 365 program.

## Conclusion

Due to the fast rate of viral mutations of SARS CoV-2 that result in decrease in the efficacy of the vaccines that have been developed before the emergence of these mutations, it is necessary to provide another approach to combat viral replication cycle. Enzymes that are required for viral replication such as polymerases, proteases, etc. present viable targets for this strategy. Thus, it is believed that using additional measures to combat the virus is not only advisable but also beneficial. The two antiviral drugs authorized for emergency use by the FDA, namely Pfizer’s two-drug regimen sold under the brand name Paxlovid, and Merck’s drug Lagevrio target proteases and induce errors in the genetic code of the virus, respectively. We believe the armament against the virus can be augmented by the addition of another class of enzyme inhibitors that are required for viral survival and its ability to replicate. Enzymes like nsp14 and nsp10/16 methyltransferases represent another class of drug targets since they are required for viral RNA translation and evading the host immune system. In this communication, we have successfully verified that the Methyltransferase Glo, which is universal and homogeneous methyltransferase assay can be used to screen for inhibitors of the two pivotal enzymes nsp14 and nsp16 of SARS CoV-2. Furthermore, we have carried out extensive studies on those enzymes using different RNA substrates and tested their activity using various inhibitors and verified the utility of this assay for use in drug screening programs. We anticipate our work will be pursued further to screen for a large library to discover new and selective inhibitors for the viral enzymes particularly that these enzymes are structurally different from their mammalian counterparts.

## Acknowledgement

The authors acknowledge Eric Yao of Signalchem Lifesciences, Richmond, BC V6V 2J2, Canada for his service in cloning and expression of nsp10, nsp16 and nsp14.

